# Coevolutionary constraints of Zika virus nonstructural protein 5 replication and interferon antagonism activities

**DOI:** 10.64898/2025.12.23.696258

**Authors:** R. Blake Richardson, Caroline Kikawa, Amit Garg, Eva Bednarski, Madihah Salim, David Bacsik, Ethan C. Veit, Anna Hermacinski, Rachael Hamilton, Adolfo García-Sastre, Jesse D. Bloom, Jean K. Lim, Matthew J. Evans

## Abstract

The flavivirus nonstructural protein 5 performs multiple functions during infection, including RNA replication and type I interferon signaling antagonism. Although flavivirus NS5 proteins inhibit IFN signaling through distinct mechanisms, which suggests evolutionary flexibility, the evolutionary constraints for these activities to coexist within a single protein remain to be determined. Here, we mapped the Zika virus NS5 STAT2 antagonism determinants and compared them with replication constraints defined by deep mutational scanning. Antagonism and replication determinant extensively overlapped, and no single amino acid substitution eliminated antagonism without impairing replication. Resolving these fitness landscapes in parallel identified specific combinations of partially functional substitutions that retained replication capacity while markedly reducing antagonism. These viruses were profoundly attenuated in human STAT2 knock-in mice. Our results uncover a fundamental evolutionary constraint linking replication and immune evasion activities in NS5, highlight that STAT2 antagonism is essential for ZIKV pathogenesis and provide new avenues for attenuated ZIKV vaccines.

## Introduction

Flaviviruses must evade host innate immune responses to establish infection, yet their replication machinery is embedded within the same proteins that counteract these responses^1,2^. This overlap presents a potential evolutionary constraint: mutations that improve immune evasion may impair replication, and vice versa. Understanding how flaviviruses manage this tradeoff is essential for defining how they adapt to immune pressures and evolve pathogenic traits. Their small genome enforces multifunctionality, which maximizes efficiency but also increases the likelihood that distinct activities are genetically and structurally intertwined. Nonstructural protein 5 (NS5), the largest flavivirus protein, exemplifies this principle because it combines enzymatic RNA dependent RNA polymerase and methyltransferase functions with antagonism of type I IFN signaling (IFN)^1^. One potential solution to this conflict would be to spatially separate determinants of IFN antagonism from the enzymatic surfaces needed for RNA synthesis and capping, which would allow each function to evolve with greater independence. To what extent IFN antagonism determinants and replication determinants overlap within NS5 remains unclear.

The NS5 proteins of different flaviviruses antagonize IFN signaling through diverse mechanisms. Some, like Zika virus (ZIKV) and DENV, target STAT2^3,4^, a key transcription factor downstream of IFNAR engagement, while others disrupt other IFN signaling nodes like TYK2 or IFNAR surface expression^5,6^. This diversity indicates a substantial evolutionary flexibility in how flaviviruses suppress antiviral signaling and is likely influenced by the cellular environments, host species, and vector relationships that define each virus’s ecological niche. Host evolution further shapes NS5 specificity and activity. ZIKV NS5-mediated antagonism is strongly species dependent. Human STAT2 is susceptible to ZIKV NS5, whereas STAT2 from *Mus musculus* is resistant^3,7–9^, which leads to rapid viral clearance and a lack of disease in immunocompetent mice^10^. In contrast, infection and pathogenesis occurs in mice when the IFN pathway is disabled or when mouse STAT2 is replaced with the human ortholog^10,11^. These observations indicate that NS5 must remain compatible with the particular innate immune signaling network of each host population. They further suggest that flaviviruses do not follow a simple linear arms race with IFN signaling components. Instead, they likely navigate a more complex evolutionary landscape where NS5 function and specificity can shift to different cellular targets when one mode of antagonism is ineffective, while still maintaining critical replication activities within the same protein.

Here, we investigate how the replication and IFN antagonism activities of ZIKV NS5 coexist. We focused on the subset of NS5 residues that physically contact STAT2 to determine how mutational flexibility within this interface contributes to antagonism. We found that the residues most critical for STAT2 antagonism also proved essential for replication, and no single amino acid substitution could eliminate antagonism without reducing viral fitness. However, by resolving both antagonism and replication landscapes in parallel, we identified mutations that in combination selectively lessened antagonism while preserving replication. These findings define a fundamental tradeoff in flavivirus evolution and demonstrate how multifunctionality within NS5 restricts adaptive flexibility, particularly when a single protein must sustain viral replication while counteracting host immunity. This work provides mechanistic insight into how these activities are linked and also suggests a strategy for rational design of immune attenuated viruses for vaccine applications.

## Results

### Surveying the genetic flexibility of ZIKV NS5 STAT2 antagonism determinants

To define the genetic requirements for ZIKV NS5 antagonism of IFN signaling, we focused on residues that directly contact STAT2^12^. Using the protein interface exploration tools PDBePISA^13^ and Protein Contact Atlas^14^, we identified 66 ZIKV NS5 amino acids that constitute an extensive interface with STAT2 (Supplementary Fig. 1, yellow residues). The majority of these contacts (49 residues) are in the RNA-dependent RNA polymerase (RdRp) domain of NS5, which interacts with the STAT2 coiled-coil domain (CCD) and amino-terminal domain (ND). An additional 17 residues in the methyltransferase (MTase) domain contact the STAT2 CCD, extending the interaction surface across both enzymatic domains of NS5. We generated 66 individual NS5 plasmid libraries, each containing randomized codons at a single STAT2-contacting residue to represent all 20 amino acids and a stop codon. We tested the ability of these libraries to inhibit IFN signaling using a 293T cell-based ISRE-luciferase reporter assay^15^. To ensure accurate determination of reporter levels in the complete absence of IFN signaling, we used STAT2 knockout 293T cells in which STAT2 complementation restores signaling (Fig 1a, compare first and second bars)^9^. Co-transfection of a wild-type (WT) ZIKV NS5 plasmid abolished IFN signaling, demonstrating its potent antagonism activity (Fig 1a, third bar). In contrast, a mutant with a substitution at a residue previously shown to be important for antagonism^12^ (D734R) failed to suppress reporter activity. We titrated WT and D734R NS5 plasmids to show that mixtures of antagonistic and non-antagonistic variants yield a graded spectrum of IFN signaling suppression (Fig. 1a). This titration curve served as a reference for interpreting the intermediate activities observed in the NS5 mutant libraries.

**Fig. 1:**
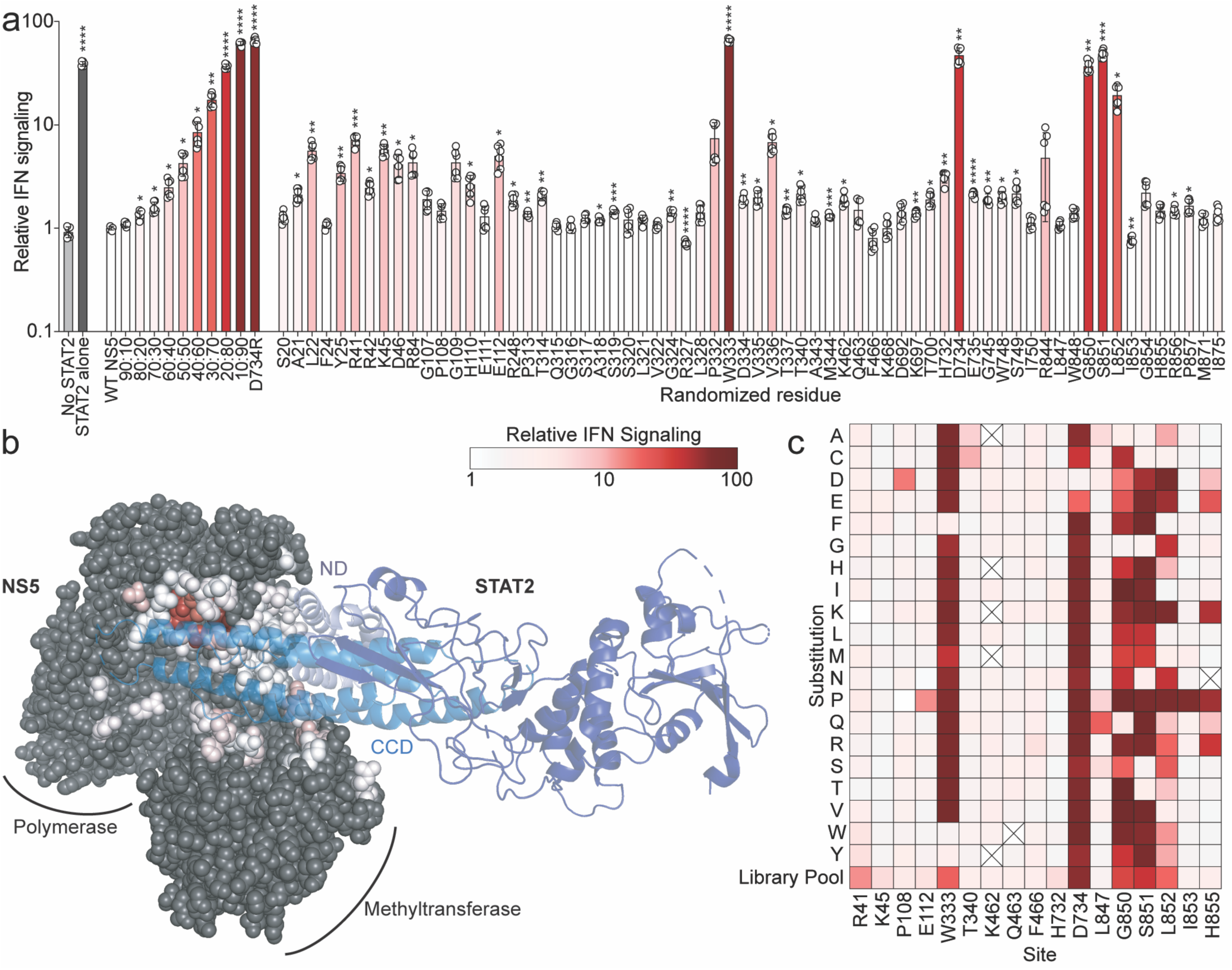
Definition of ZIKV NS5 STAT2 antagonism determinant genetic flexibility. **a** Relative IFN signaling (mean ± s.d., *n* = 6), defined as the ratio of constitutively expressed Renilla luciferase to ISRE-driven firefly luciferase activity, measured in STAT2 knockout 293T cells. The first two bars demonstrate reporter ratios in the absence of IFN signaling (‘No STAT2’, light gray) and co-transfected STAT2 plasmid (dark gray). All remaining bars include ZIKV NS5 expression and are color-coded to indicate the level of IFN signaling (white = low signaling/strong antagonist; dark red = high signaling/inactive antagonist), using the scale shown below and consistent across the figure. The first NS5 condition is a standard curve composed of defined ratios of wild-type (active antagonist) and D734R (inactive antagonist) NS5 plasmids, demonstrating the assay’s capacity to resolve intermediate phenotypes. ‘Randomized residue’ bars represent mixtures of NS5 plasmids encoding randomized single amino acid substitutions at the indicated residue (x-axis). Asterisks indicate statistical significance determined by multiple unpaired t-tests with Holm-Šidák method to correct for multiple comparisons. **b** Structure of the ZIKV NS5–STAT2 complex (composite of PDB: 6WCZ, 6UX2), showing NS5 in space-filling representation and STAT2 in ribbon format. NS5 residues in contact with STAT2 are heat-mapped according to the relative IFN antagonism mutational sensitivity measured in panel *a*. **c** Heatmap showing the impact of the indicated substitution (rows) at each NS5 site (columns) on IFN signaling levels. Each square is colored using the same scale as in panels *a* and *b*, with darker red indicating higher IFN signaling. Squares with X’s indicate mutants that were not tested. Primary data for individual mutants are shown in Supplementary Fig. 2. All IFN signaling values in this, and subsequent figures represent means plus and minus standard deviation from at least two independent experiments each with three technical replicates.

Antagonism activity varied substantially among NS5 plasmid libraries. Most residues tolerated substitutions, with their libraries retaining near–wild-type antagonism activity (Fig. 1a, white bars). Several libraries, however, exhibited weak to moderate reductions in antagonism (Fig. 1a, pink bars). Because each library contains all possible substitutions at a single site, the observed effect reflects the combined impact of those variants. Moderate decreases in activity could result from mild impairments across many substitutions or from strong defects caused by a subset of particularly disruptive mutants. Four residues (W333, D734, G850, and S851) were generally intolerant of mutation, as their corresponding libraries showed a near-complete loss of antagonism activity (Fig. 1a, dark red bars). Mapping the relative antagonism capacity of each mutant library onto the NS5 structure revealed that the most sensitive residues cluster in the thumb region of the NS5 RdRp domain, where they contact the distal end of the α-helices forming the STAT2 CCD (Fig. 1b).

To validate the above survey and further define the mutational tolerance of NS5 antagonism determinants, we tested the antagonism activity individual amino acid substitutions at a subset of STAT2 contact residues (Fig. 1c and Supplementary Fig. 2). Residues that appeared highly tolerant to mutations in the randomized mutant libraries (Fig. 1a) exhibited similar tolerance of individual amino acid substitutions (Fig. 1c and Supplementary Fig. 2a-d, f-j, l, and p). Conversely, at residues where the random mutant libraries exhibited greatly decreased antagonism (Fig. 1a), the majority of individual single amino acid mutants also impaired antagonism (Fig. 1c and Supplementary Fig. 2e, k, m, n, and o). Thus, we defined the relative mutational tolerance of the NS5–STAT2 interface and identified a discrete structural feature that is critical for IFN antagonism.

### Comparing ZIKV NS5 STAT2 antagonism determinants across flaviviruses

We next sought to determine whether residues critical for STAT2 antagonism in ZIKV NS5 also govern antagonism in other flaviviruses. Two mutations, D734R and S851R, abolished ZIKV NS5’s IFN signaling antagonism (Fig. 1c, Supplementary Fig. 2k and n, and Fig. 2a). Introducing equivalent substitutions into SPOV, DENV2, and YFV NS5 similarly eliminated their antagonism activity (Fig. 2a, dark orange, light orange, and yellow bars, respectively). Since these viruses target STAT2, the shared sensitivity to mutation suggests a conserved STAT2-binding interface. In contrast, the same substitutions had minimal impact on activity on IFN antagonism by LGTV and WNV NS5, which target TYK2^5^ and PEPD^6^, respectively, which were tested in parallel with known antagonism defective mutants of each (LGTV NS5 with the YFV variable loop (VR)^5^ and a naturally occurring F653S mutation in the attenuated WNV Kunjin strain^16^) (Fig. 2a, green and blue bars, respectively). Importantly, differences in antagonism did not correlate with loss of NS5 expression (Fig. 2b). Together, these results support a model in which closely related flaviviruses share a conserved STAT2-targeting interface on NS5, whereas more divergent species that target TYK2, PEPD, or other IFN signaling components rely on distinct molecular surfaces.

**Fig. 2:**
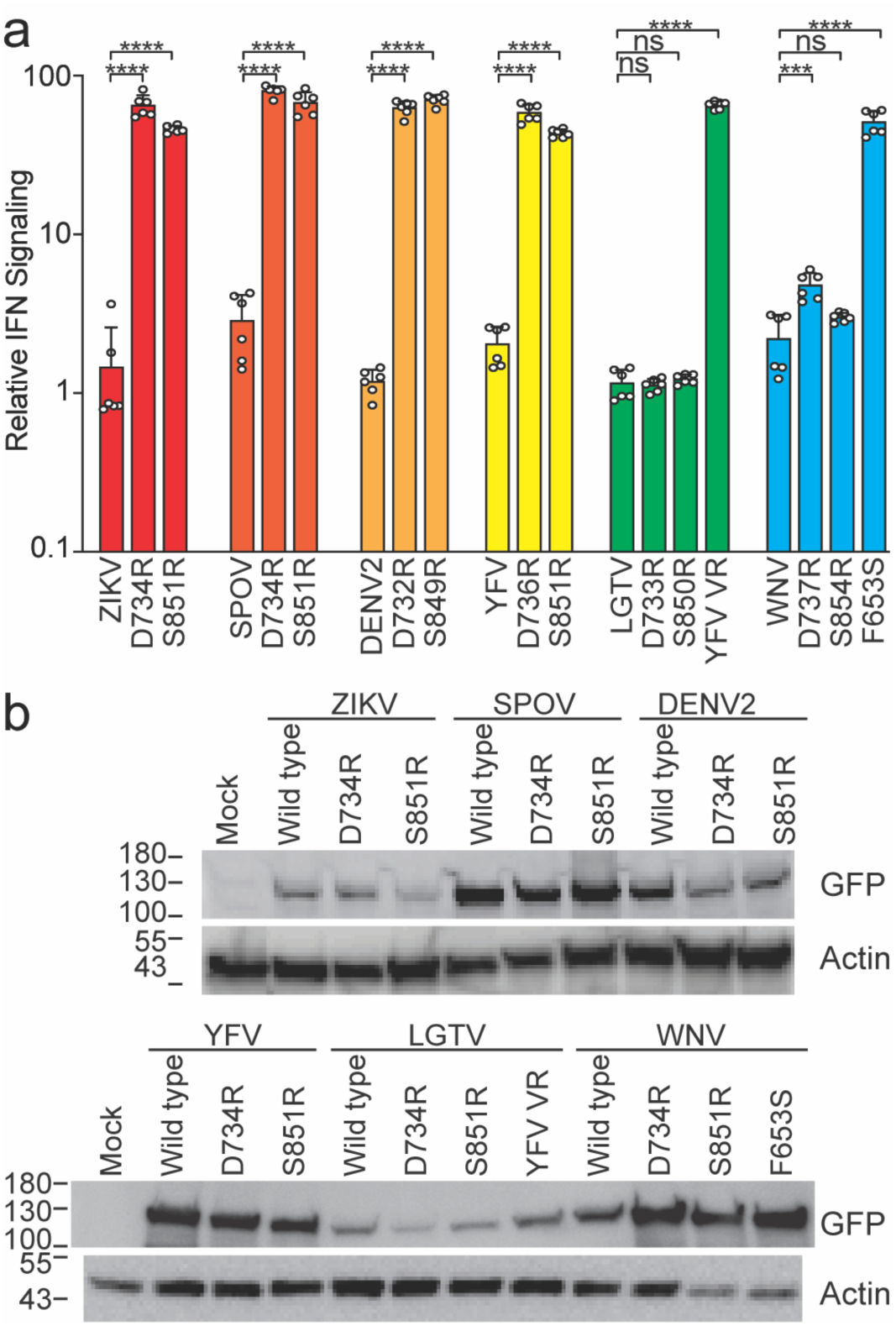
Conservation of STAT2 antagonism determinants between flaviviruses. **a** Relative IFN signaling levels in 293T cells transfected with the indicated flavivirus NS5 expression plasmid. Wild type and mutant versions from the same virus are color-coded. Asterisks indicate statistical significance determined by unpaired t-test with Welch’s correction. **b** Immunoblots of lysates from the above transfections with an antibody that binds GFP, which is fused to the amino terminus of all the NS5 proteins, or actin, as a loading control. Molecular weight marker (kDa) positions are labeled on the left.

### Defining ZIKV NS5 replication determinants

To map the genetic constraints imposed on ZIKV NS5 by its role in replication and to compare these with those required for IFN antagonism, we performed deep mutational scanning (DMS) in the context of a live virus. This approach involves serially passaging pooled viral libraries containing all possible single amino acid substitutions in NS5, allowing the selection of variants that support efficient viral replication. Mutations that are depleted over time likely disrupt replication, while enriched variants are compatible with the viral life cycle.

Mutant virus pools were passaged through Huh-7.5 cells, which lack a functional RIG-I mediated antiviral response, thereby eliminating interferon signaling as a selective pressure and ensuring that observed differences in enrichment reflect replication capacity rather than immune antagonism^17^. Infections were performed at a multiplicity of infection (MOI) of 0.05 infectious units per cell to ensure that each were initiated by single viral particles, establishing a genotype:phenotype linkage and allowing individual mutant genomes to at least initially replicate and spread independently. Two days after infection, when the monolayer was nearly fully infected, we harvested total cellular RNA. At this stage, variants that had successfully completed multiple rounds of replication and spread should be enriched in the population, while those with replication defects should be depleted.

We used Illumina-based next generation sequencing and a previously described barcoded-subamplicon error correction method^18,19^ to show that most possible mutations are sampled very well in the plasmid samples and the proportion of mutants drops substantially following selection in Huh-7.5 cells (Supplementary Fig. 3a and b). Furthermore, stop codon substitution mutants were purged following passaging, indicating effective selection for fit mutants (Supplementary Fig. 3a). We compared the relative proportions of mutants in the pre- and post-selection samples to estimate the impact of each substitution on viral replication in Huh-7.5 cells, which we term ‘mutational effect’. These values represent enrichments of each amino acid at a site after selection for viral growth, normalized to the abundance of the wild-type codon at each site (Supplemental Table 1, top of ‘DMS Results’ tab). The observed mutational effects strongly correlated among three biological replicate libraries (Supplementary Fig. 3c), indicating reproducible mutant selection. Hereafter, we report the median across the three replicate libraries.

Figure 3a shows the cumulative mutational effect of all single amino acid mutations across all of ZIKV NS5. In this heatmap, mutants that have impaired replication in Huh-7.5 cells relative to the WT amino acid at that site are shades of blue to deep purple, while mutants that had an increased replication over the WT amino acid are bright green to yellow. Amino acids known to be involved in NS5 methyltransferase and polymerase activities tolerated few to no amino acid substitutions (e.g., K61, D146, K182 and E218 for the methyltransferase active site, G664-D666 for the polymerase active site) (Fig. 3c). This further supports that the DMS screen correctly measured individual mutant effects on viral replication. To validate our data, we generated a subset of single amino acid NS5 mutants in a ZIKV cDNA clone encoding a luciferase reporter gene. We rescued stocks of these viruses by transfection of 293T cells and quantified their replication by infecting Huh-7.5 cells and scoring luciferase activity 2 days later. In general, luciferase activity in infected cells closely matched the effects of each mutant measured in deep mutational scanning (Fig. 3b). Specifically, substitutions with a DMS predicted mutational effect of 0.1 or less (e.g.10-fold lower than the endogenous residue) strongly impaired viral growth, whereas mutations that had wild-type or better scores in the deep mutational scanning always grew well in the growth curves, even if these mutations are not observed among natural sequences. However, mutations that appeared to be just mildly attenuated in the deep mutational scanning (selected at a frequency within 10-fold of the wild-type frequency) often grew about as well as the wild type, suggesting that caution should be used in calling mutations deleterious if they were only mildly depleted in the high-throughput experiments.

**Fig. 3:**
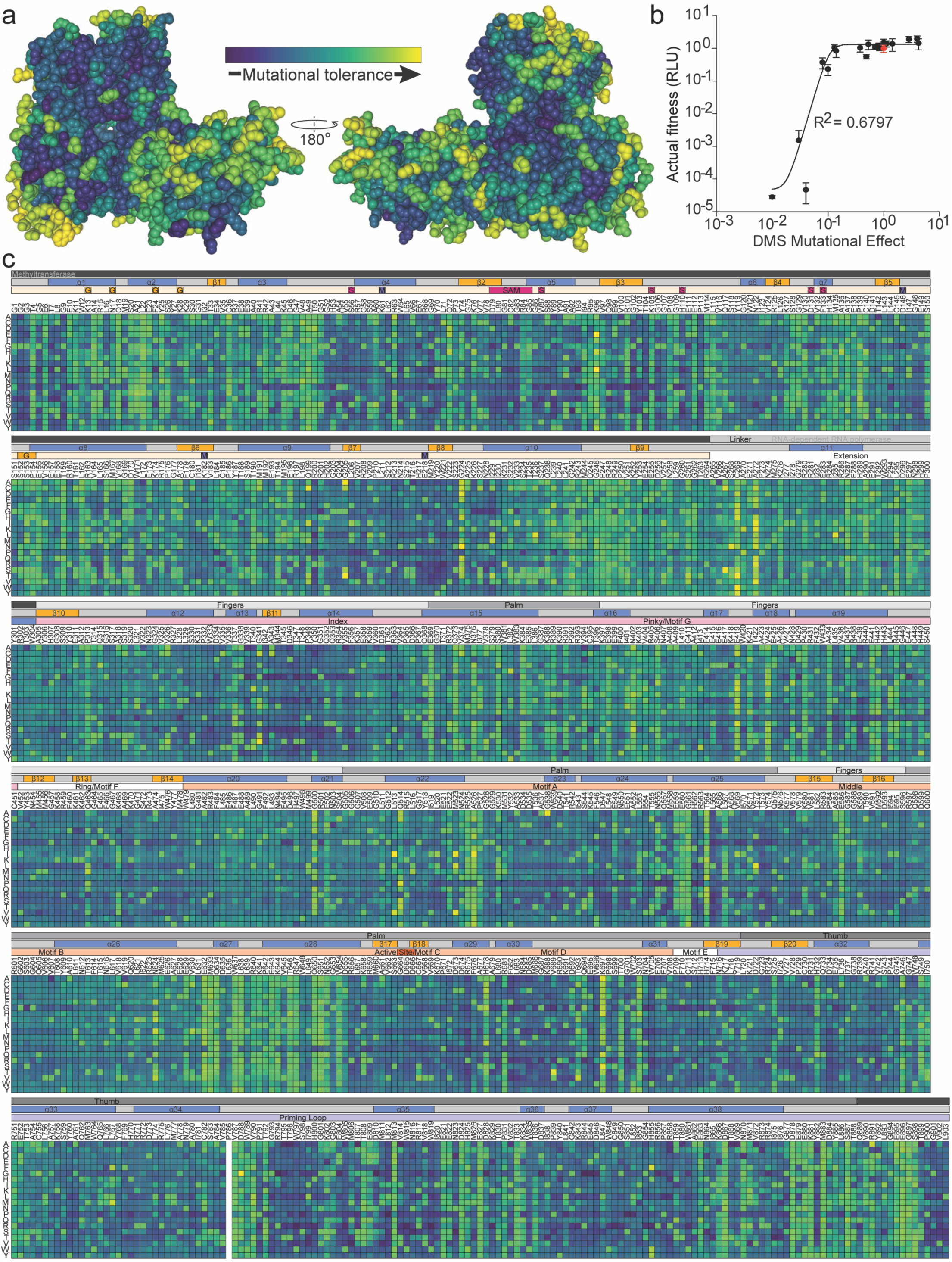
Deep mutational scanning of ZIKV NS5 replication determinants in Huh-7.5 cells. **a** Experimentally measured mutational tolerance mapped onto the ZIKV NS5 structure (composite of PDB: 5TFR and 5U04). Mutational tolerance at each site is quantified as the median enrichment effect of all single amino acid substitutions across three biological replicates, such that sites that tolerate many amino acids have high mutational tolerance and sites that strongly prefer the wild-type residue have low mutational tolerance. This tolerance metric was then used to color the NS5 structure (yellow = high tolerance; purple = strong constraint). Two views rotated 180° are shown. **b** Heatmap of mutational effects for every possible single amino acid substitution across NS5 (columns = sites, rows = amino acid identities). White squares indicate substitutions whose effects could not be confidently resolved. Annotations above the heatmap (top to bottom) indicate: NS5 domain organization (MTase and RdRp), secondary structural elements (α = α-helix, β = β-strand), and key functional features (G = 7-MeGpp interaction site, S = SAM-binding region, M = K-D-K-E methyltransferase catalytic tetrad), inferred from^35^. White squares indicate substitutions whose effects could not be confidently resolved. Values represent the median mutational effect across three independently generated tile libraries for each NS5 segment, normalized to wild-type residue preference.

### Comparing ZIKV NS5 STAT2 antagonism and replication genetic constraints

To determine the degree of overlap between antagonism activity and replication determinants of NS5 residues, we compared the data from our antagonism experiments with our DMS experiment. Across the entire NS5 protein, our DMS results predict that an average of 7.0 amino acids substitutions would be tolerated for replication (defined as substitutions that exhibit a mutational effect of great than 0.1). In general, residues in the STAT2 interface had a similar replication mutational tolerance of 8.3 different amino acids at each position. However, the residues deemed genetically constrained for STAT2 antagonism (W333, D734, G850, and S851) (Fig 1a) exhibited a replication tolerance of only 1.5 different amino acids. Indeed, only the WT residue is predicted to function for replication for W333 and G850, while D734 can also tolerate an E substitution and S851 could tolerate an A substitution (mutational effect of 0.10 and 0.20, respectively). The correlation of these activities suggests that the same residues required for antagonism are also required for replication. Specifically, the residues that are the most constrained for antagonism also show a reduced mutational tolerance when measuring replication fitness. This suggests these key amino acids are under a selective pressure to maintain dual functionalities.

### Separating ZIKV NS5 IFN antagonism and replication functions

Disconnecting the replication and IFN antagonism functions of NS5 could enhance our understanding of viral evolution under host immune pressures, provide valuable insights into viral pathogenesis and immune evasion strategies, and facilitate the development of live attenuated vaccines. However, the above comparison of IFN antagonism and replication genetic constraints suggests this might be difficult. By plotting the 293T cell IFN signaling assay results and the DMS predicted fitness values of 435 individual mutants tested at 62 different NS5 STAT2 contact sites, we again found a strong correlation between these activities (Fig. 4a). Many mutants exhibited strong IFN antagonism, as gauged by a low y-axis ‘Relative IFN signaling’ result, and strong predicted viral replication capacity, as reflected by a DMS mutational effect greater than 0.1 (Fig. 4a, lower right quadrant). Many other mutants were impaired for both functions (Fig. 4a, top left quadrant) or capable of antagonizing IFN signaling but predicted to be highly impaired for replication (Fig. 4a, bottom left quadrant).

**Fig. 4:**
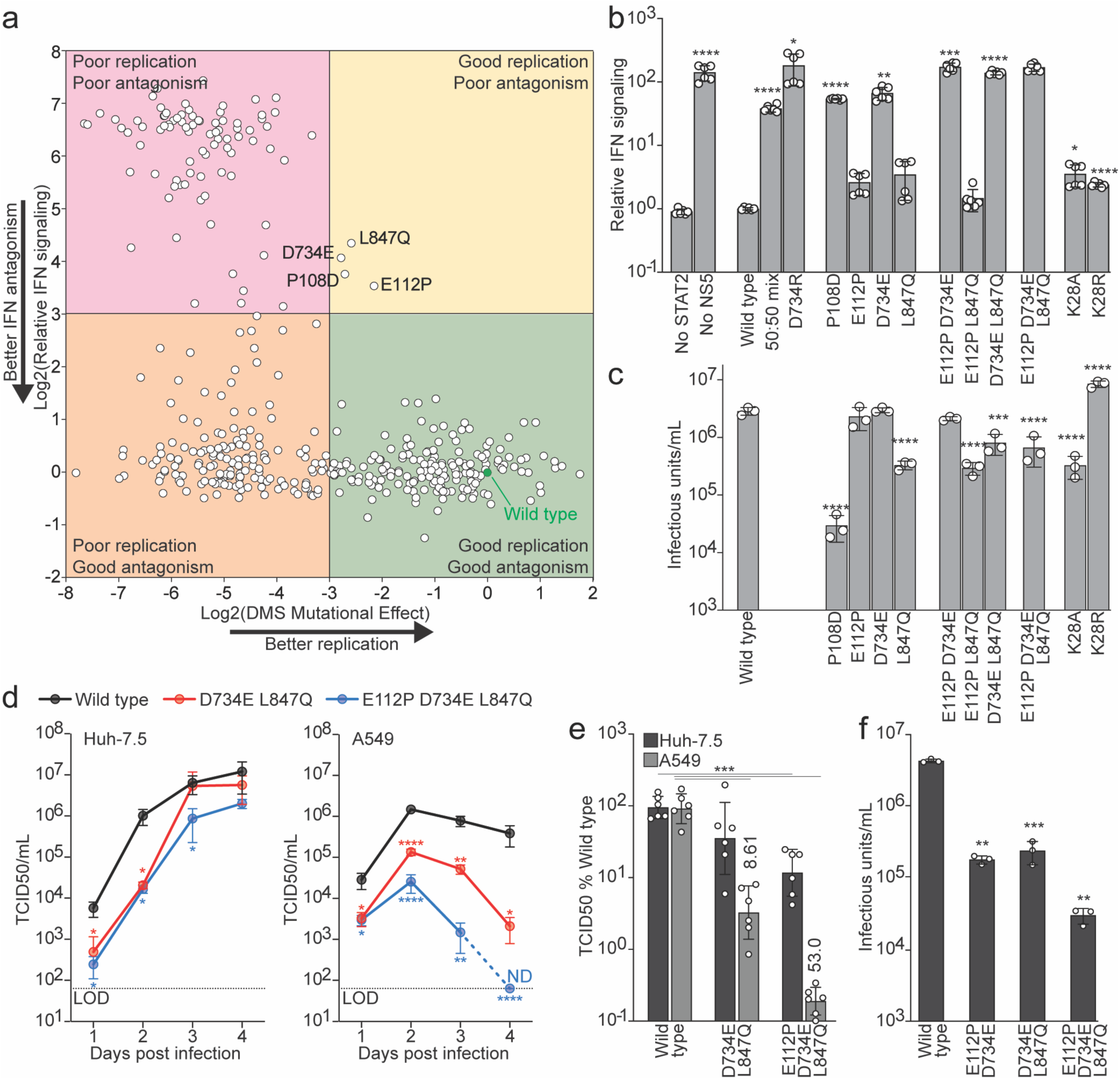
Separating ZIKV NS5 IFN antagonism and replication functions. **a** Comparison of replication fitness and STAT2 antagonism effects for all NS5 substitutions identified by deep mutational scanning and single-mutant antagonism assays. Each point represents a single amino acid substitution plotted by its Huh-7.5 cell DMS predicted replication mutational effect (x-axis) and impact on 293T cell IFN signaling (y-axis) and normalized to the wild type NS5 values (green filled circle). Quadrants indicate phenotypic classes (for instance lower right = good replication and antagonism). Mutants evaluated in downstream assays are labeled. **b** STAT2 antagonism activity of individual and combinatorial NS5 mutants measured by luciferase reporter assay in HEK293T cells treated with IFN. Higher values indicate greater weaker antagonism. Asterisks indicate statistical significance determined by multiple unpaired t-tests with Holm-Šidák method to correct for multiple comparisons. **c** Replication fitness of ZIKV encoding the same substitution mutants, quantified as infectious virus produced in from transfection of 293T cells transfected with plasmids encoding each virus genome and quantified by E protein FACS on Huh-7.5 cells. Asterisks indicate statistical significance determined by multiple unpaired t-tests. **d** Growth kinetics of selected mutants in IFN-deficient Huh-7.5 cells and IFN-competent A549 cells, quantified by TCID50/mL on STAT2 KO Huh-7.5 cells at indicated days post-infection. Dotted lines denotes assay limits of detection (LOD) and ND = not detected. Asterisks indicate statistical significance determined by multiple unpaired t-tests. **e** Three day post infection titers from panel d expressed as a percentage of wild-type levels in each cell line, highlighting diminished replication specifically in the IFN-competent background. Numbers indicate the fold difference between Huh-7.5 and A549 cell titers. Asterisks indicate statistical significance determined by multiple unpaired t-tests. **f** Infectious titers of mouse-adapted wild type and mutant viruses for in vivo experiments produced from a Dakar-41525 infectious clone produced from 293T cells and tittered by E protein FACS on Huh-7.5 cells.

**Fig. 5:**
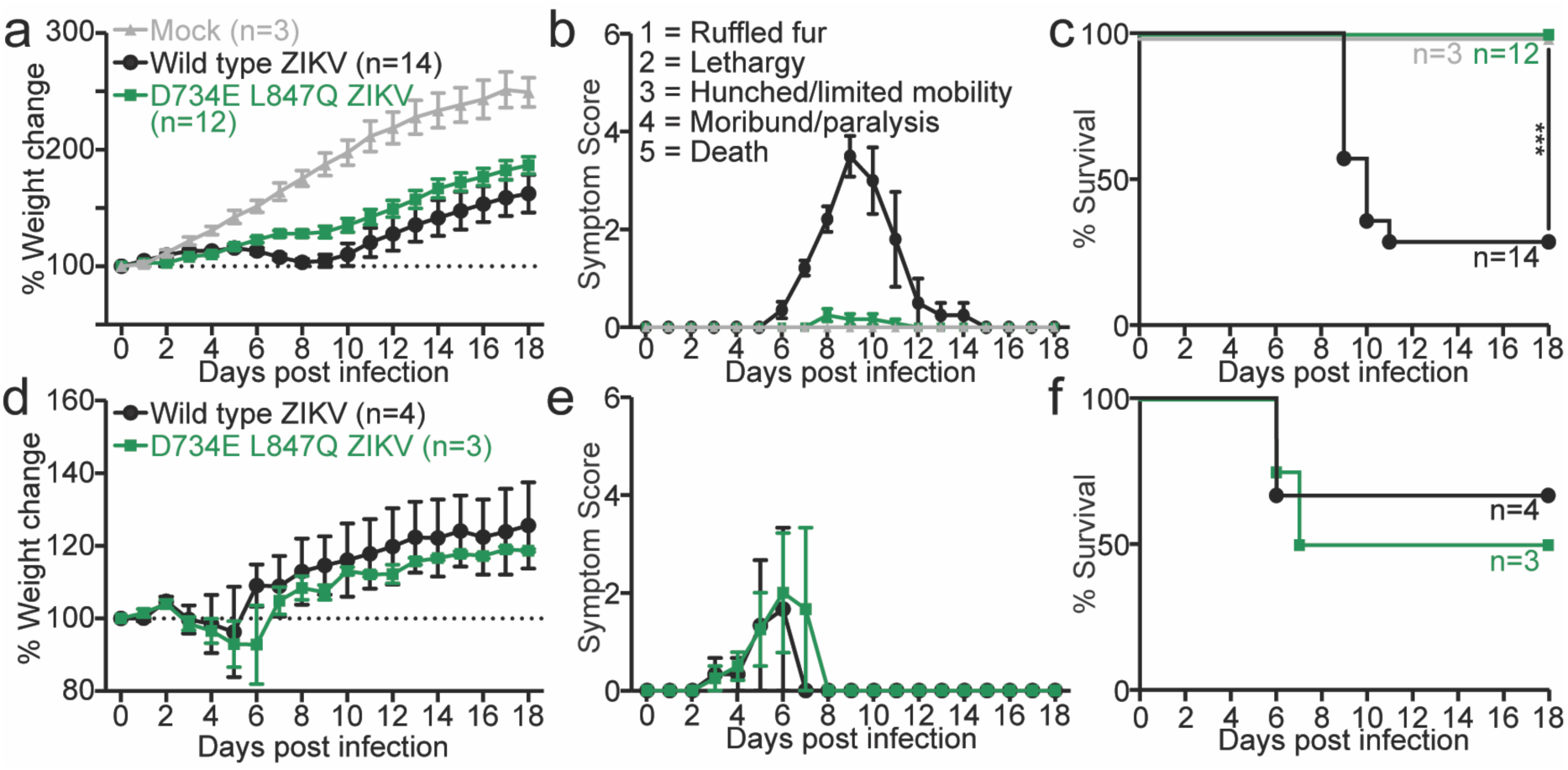
Loss of NS5-dependent STAT2 antagonism severely attenuates ZIKV disease in vivo. Human STAT2 knock-in (*hSTAT2^KI/KI^*) mice (**a–c**) or STAT2-deficient (*Stat2-/-*) mice (**d–f**) were infected with equal infectious doses of wild-type (WT) or STAT2-antagonism-deficient D734E/L847Q ZIKV. In *hSTAT2^KI/KI^* mice, WT infection led to sustained weight loss (**a**), neurological disease (**b**), and 60% mortality (**c**), whereas D734E/L847Q infection caused minimal symptoms and no mortality despite detectable replication (viral RNA in brain, spinal cord, and spleen at 7 dpi). In *Stat2-/-* mice, both viruses caused similar weight loss (**d**), symptom progression (**e**), and mortality (**f**), indicating that attenuation of the mutant virus results specifically from its inability to antagonize STAT2. Data represent pooled biological replicates with mean ± s.e.m.; survival assessed by log-rank test.

To identify variants that reduce IFN antagonism while preserving viral replication, we examined NS5 substitutions that clustered in the top right quadrant in Figure 4a. These variants showed a measurable loss of antagonism, yet their deep mutational scanning effects remained above the 0.1-fold cutoff that we used to define replication incompatibility. Four mutants met this criterion: P108D, E112P, D734E, and L847Q. We first evaluated each mutant individually by measuring STAT2 antagonism side by side with wild-type NS5 (Figure 4b). All four substitutions impaired antagonism, although to varying extents. Certain combinations further reduced antagonism activity, and a subset performed nearly as poorly as the no-NS5 control. We then tested the ability of each mutant to support viral replication by rescuing infectious virus and determining titers in cell culture (Figure 4c). P108D caused an approximately 80-fold reduction in viral yield, so we excluded this variant from further study. The remaining mutations were broadly tolerated. E112P and D734E supported replication at levels similar to wild type, and L847Q resulted in a ten-fold reduction.

We next tested combinations of the three well tolerated substitutions. Several double mutant viruses exhibited little or no additional loss of fitness relative to the single mutant controls (Figure 4c). Based on these results, we selected the D734E L847Q double mutant and the E112P D734E L847Q triple mutant for further characterization. These variants showed the strongest reduction in antagonism while maintaining sufficient replication to support downstream studies in more physiologically relevant settings.

A recent study identified two ZIKV NS5 mutations, K28A and K28R, at a site near, but not directly interacting with STAT2, that selectively reduce STAT2 antagonism while preserving replication^20^. To compare those findings with our broader analysis, we tested K28A and K28R in our assays. As previously reported, both K28A and K28R mutants modestly reduced antagonism activity (Fig. 4b) while maintaining robust replication, with K28R even producing slightly higher titers than wild type (Fig. 4c). These results are consistent with our finding that single amino acid NS5 changes are limited in their ability to strongly alter antagonism without incurring replication costs, underscoring the significance of the multi-step mutational trajectories required to functionally separate these two activities.

We next evaluated multicycle replication of the D734E L847Q double mutant and E112P D734E L847Q triple mutant in two cell types that differ in their innate immune signaling capacity. Again, Huh-7.5 cells lack a functional RIG-I pathway and therefore support viral replication without innate immune restriction, whereas A549 cells mount a robust IFN response. Each virus was used to infect both cell types and supernatants were collected over time. Infectious titers were quantified by TCID50 (Figure 4d). Mutants with impaired STAT2 antagonism showed disproportionately reduced replication in A549 cells compared with Huh-7.5 cells, consistent with a specific defect in countering innate immunity rather than a general loss of fitness. To further quantify this immune-dependent replication difference, we expressed titers from each cell line as a percentage of wild type levels at 3 days post infection (Figure 4e). At this time point, the D734E L847Q double mutant titer produced 59% of wild type virus titers from Huh-7.5 cells, but only 4.2% of wild virus titers from A549 cells. The E112P D734E L847Q triple mutant showed a greater defect, with 14% and 0.21% of wild type titers from Huh-7.5 and A549 cells, respectively. These results confirm that the selected NS5 variants retain the capacity to replicate, but have a significantly diminished ability to suppress antiviral signaling in cells with an intact IFN response.

### Loss of IFN antagonism attenuates ZIKV *in vivo*

To test the in vivo relevance of NS5 mediated STAT2 antagonism, we generated an infectious cDNA clone of the mouse adapted Dakar-41525 strain bearing the NS4B G18R mutation that enhances replication in mice, termed ZIKV-Dak-MA^21^. We rescued wild-type and mutant NS5 ZIKV-Dak-MA viruses and quantified infectious titers in cell culture (Figure 4f). The D734E L847Q double mutant replicated to titers that were about one order of magnitude lower than WT, indicating that its replication capacity remained largely intact. The E112P D734E L847Q triple mutant displayed a larger decrease in yield but still produced readily detectable infectious virus. These findings show that the mutations that impair STAT2 antagonism remain compatible with viral replication, allowing us to assess their functional consequences in vivo.

To determine whether impaired STAT2 antagonism limits disease *in vivo*, we infected human STAT2 knock-in mice (*hSTAT2^KI/KI^*) with equivalent doses of wild-type virus or the D734E L847Q double mutant. All *hSTAT2^KI/KI^* mice infected with wild-type virus displayed increase indications of disease as measured by greater weight loss, with all mice developing clinical symptoms compared to the mice infected with the D734E L847Q double mutants, which showed less weight loss, with mice showing either no symptoms or mild symptoms. Consistent with these data, the mice infected with the wild type virus had significantly lower survival rates (28%) compared to these same mice infected with the double mutant, where all mice survived (p=0.0003). To confirm that the mutant virus retains the capacity to cause disease when innate immune restriction is removed, we infected Stat2 knockout mice (*Stat2-/-*) with lower inoculum of WT and double mutant virus. In this background, both wild-type and mutant viruses caused a similar level of weight loss, developed similar symptoms, and survival was comparable. These data demonstrate that NS5 mediated STAT2 antagonism is required for ZIKV pathogenesis *in vivo* and that selective disruption of this function produces a strongly attenuated phenotype.

## Discussion

Flaviviruses must simultaneously efficiently replicate their genomes and evade innate immunity, yet these demands often encoded within the same multifunctional NS5 protein. How these distinct activities coexist and coevolve within this protein has remained unclear. Here, we reveal that the ZIKV NS5 STAT2-binding footprint highly restricted and embedded within a region under strong replication constraint. Mutational profiling showed that residues critical for antagonism uniformly overlap with sites required for robust viral fitness, indicating that STAT2 antagonism likely evolved atop pre-existing replication determinants. As a result, there are limited mutational paths that selectively disrupt antagonism without jeopardizing replication. We speculate that viruses closely related to ZIKV, such as SPOV, DENV, and YFV, would have similar evolutionary constraint on their residues since the interface between STAT2 and the respective NS5 of these species is similar. Recently the same key residues have been shown to be involved in this interface for several of these viruses^22^.

This tight linkage suggests that NS5 evolution proceeded through stepwise acquisition of STAT2 antagonism-enhancing mutations paired with compensatory changes that preserve replication. Such a trajectory would allow adaptation to new host environments while avoiding catastrophic loss of fitness. The same architecture may now constrain future evolutionary flexibility: when STAT2 antagonism is suboptimal in a given host species, it may be easier for a virus to evolve a distinct immune evasion strategy than to extensively remodel STAT2 antagonism determinants in a replication competent manner. For instance, we have recently shown that STAT2 antagonism is a barrier to ZIKV infection of rodents^9,21,23^. It is unlikely that changes that enhance mouse STAT2 antagonism would be permissive to replication. As tick-borne and WNV-related flaviviruses antagonize TYK2 and PEPD with determinants that appear distinct from STAT2 interfaces, it is possible that the antagonism and replication functions are more separated for these viruses.

Nonetheless, specific combinatorial substitutions can partially decouple ZIKV NS5 replication and antagonism functions. The D734E and L847Q double mutant retained sufficient replication capacity but was rendered largely unable to suppress STAT2-dependent signaling. This separation of function reveals the direct contribution of antagonism to disease. In human STAT2 knock-in mice, wild-type ZIKV caused severe disease and mortality, whereas the D734E L847Q virus, despite replicating in vivo, was profoundly attenuated and completely non-lethal. Importantly, attenuation was reversed in STAT2-deficient mice, establishing that loss of STAT2 antagonism, rather than reduced replicative capacity, drives the avirulent phenotype. These results demonstrate that innate immune antagonism is not merely a virulence enhancer but an essential component of ZIKV pathogenesis.

Finally, our findings provide a rational foundation for attenuation strategies. A virus that replicates but cannot antagonize STAT2 is intrinsically constrained by host immunity. The D734E L847Q mutant exhibits exactly this property and requires multiple nucleotide changes to revert to a fully pathogenic genotype, offering a strong genetic barrier to escape. Such “STAT2-blind” viruses represent promising candidates for live-attenuated ZIKV vaccines, and analogous strategies may be broadly generalizable to other flaviviruses whose NS5 proteins integrate replication and immune repression.

## Methods

### Cell lines and culture

293T and Huh-7.5 cells (provided by Charles M. Rice, Rockefeller University, New York, NY) were grown as previously described^24^ in Dulbecco’s Modified Eagle’s Medium (DMEM; Gibco BRL Life Technologies, Gaithersburg, MD) with 10% fetal bovine serum (FBS; Gibco BRL Life Technologies, Gaithersburg, MD), 108.8 units/mL penicillin, and 108.8 µg/mL streptomycin (Gibco BRL Life Technologies, Gaithersburg, MD). STAT2 KO HEK-293T cells have been previously described ^9^.

### Flow cytometry

Cells were fixed with 1% paraformaldehyde and stained with the primary 4G2 flavivirus E antibody (kind gift from Florian Krammer, Icahn School of Medicine at Mount Sinai, New York, NY) and a goat anti-mouse secondary antibody conjugated to Alexa fluorophore 647 (ThermoFisher Scientific, Waltham, MA), using methods previously described (Veit et al., 2024). Cells were analyzed using an Attune NxY flow cytometer (ThermoFisher Scientific, Waltham, MA).

### Expression plasmid cloning

All STAT2 and flavivirus NS5 expression vectors were generated by cloning into a modified version of pSCRPSY (GenBank accession no. KT368137.1) (a gift from Paul Bieniasz, Rockefeller University, New York City, NY), termed pSCRPSYneo, which encodes a neomycin resistance cassette in place of PAC2ATagRFP sequence. While this is a lentiviral plasmid suitable for generating a provirus for incorporation into lentiviral pseudotyped virions, we use this plasmid as an expression vector in transient transfection experiments. The pSCRPSYneo-STAT2-GFP plasmid encoding human STAT2 translationally fused at its C-terminus to GFP was previously described^9^. pSCRPSYneo-GFP-flavivirus NS5 plasmids encoding the 1947 MR766 ZIKV, SPOV SA Ar94, and DENV2 16681 NS5 sequences cloned as a translational fusion to GFP at its N-terminus and have also been previously described^9^. pSCRPSYneo-GFP-LGTV and WNV NS5 plasmids were generated by replacing the ZIKV NS5 sequence of pSCRPSYneo-GFP-ZIKV MR766 NS5 with the NS5 sequences of TP21 strain LGTV and NY99 WNV NS5, respectively, using the In-Fusion Cloning Kit (Takara Bio USA, Inc., San Jose, CA) (plasmid and oligo sequences available upon request).

### Cloning of mutant NS5 libraries

To create ZIKV NS5 libraries with each of the 66 ZIKV NS5 residues that contact STAT2 sites randomized, a single site variant library was synthesized (TWIST Bioscences, South San Francisco, CA) that was comprised of an arrayed library of 66 wells, where each well represented a pool of all possible amino acid variants at each of the STAT2 contact sites. Fragments encompassed the entire ZIKV NS5 coding sequence with 5’ and 3’ ends needed for InFusion cloning into the BsrGI and XhoI sites, respectively, of pSCRPSYneo-GFP-ZIKV MR766 NS5. InFusion reactions were transformed into Stellar competent cells (Takara Bio USA, Inc., San Jose, CA) and plated on 15 cm LB plates with 100 ug/mL ampicillin, to select for the pSCRPSY plasmid. Each transformation was performed in a sufficient scale to generate >5500 colonies, to overrepresent the 20 possible codon mutants by >260-fold, which were pooled by scraping and subjected to plasmid DNA preparation using the HiSpeed Plasmid Mini Kit (Qiagen, Germantown, MD).

A dilution of each bacterial transformation was plated to facilitate the determination of the library colony numbers and to pick individual colonies for miniprep analysis. Sanger sequencing of at least a dozen colonies for each STAT2 contact site library demonstrated that all clones encoding full length NS5 encoded a mutant codon at the intended position. A proportion of these colonies bore NS5 sequences with internal deletions of the NS5 coding sequence, which was likely caused by stretches of repetitive sequence. The frequency of full length NS5 within each library was scored by subjecting each plasmid library pool to whole plasmid sequencing (Psomagen, Rockville, MD). The proportion of raw reads that bore the entire NS5 sequence within each pool was calculated and used to correct the observed IFN signaling antagonism activity of this plasmid mixture (see Datafile 1 for analysis details).

### Construction of mouse adapted Dakar ZIKV cDNA clone

To enable genetic manipulation of a ZIKV strain capable of infecting and causing disease in humanized STAT2 mice, we generated a plasmid encoding the full-length cDNA of the Dakar-41525 strain carrying the NS4B G18R substitution, a mutation previously shown to enhance replication in mice^21^. This clone, termed ZIKV-Dak-MA, places the viral cDNA under the control of the CMV promoter and includes a hepatitis delta virus ribozyme (HDVr) followed by an SV40 polyadenylation signal at the 3′ end to generate authentic viral RNA termini, analogous to our previously described ZIKV infectious clone28. Because flavivirus sequences are toxic in bacteria, two synthetic introns were inserted after viral genome nucleotides 3126 and 8517 at regions empirically identified as unstable during cloning. Table 2 lists the DNA fragments and the cloning strategy used to assemble the final pCDNA6.2 Dakar 2Intron NS4B G18R HDVr polyA plasmid. The complete sequence of this construct has been deposited in GenBank under accession number PX701121.

### IFN signaling antagonism assay

To quantify IFN signaling and antagonism, we used a modified version of a luciferase reporter-based 293T cell assay^25,26^. One day before transfection, 50,000 STAT2 KO 293T cells were seeded in 500 uL of DMEM with 3% FBS in a 24-well plate. The next day, each well was transfected with a pool of plasmids: TK-Rluc (40 ng; constitutively expressed reporter used as transfection control), ISG54-ISRE-Fluc (200 ng; reporter induced by IFN treatment), STAT2 (3.7 ng; to complement STAT2 KO, omitted to demonstrate background reporter levels), and either pSCRPSYneo-GFP (33 ng; used as an empty vector control to illustrate full IFN signaling levels;) or pSCRPSYneo-GFP-flavivirus NS5 (33 ng; NS5 genotype is the major difference between experiments). At 24 h post-transfection, the cells with treated with 100 U of human IFNb (at 100 U/ml). Cells were lysed 24 h after IFNb treatment, and FLuc and RLuc expression was measured using the Dual-Luciferase Reporter Assay System (Promega, Madison, WI) and a BioTek Synergy H1 Multimode Reader (BioTek, Winooski, VT). Data were calculated as a ratio of Fluc:RLuc to normalize for transfection efficiency. At least two independent experiments with three biological replicates were performed for all reported values. To assay the antagonism activity of mixtures of NS5 mutants shown in Fig 1a, 33 ng of defined mixtures of wild type and D734R ZIKV or each NS5 random codon pooled library DNA were transfected.

### ZIKV NS5 DMS

To generate mutant plasmid libraries, we created ZIKV NS5 codon-mutant DNA fragments using a previously described PCR mutagenesis approach^27^ with sets of forward and reverse oligos that randomized each codon with an NNS sequence where N is any nucleotide and S is either a G or a C (Table 3). Forward and reverse oligos for each tile were synthesized as oPools Oligo Pools (Integrated DNA Technologies, Coralville, IA). The sequences of these primers are included in Table 3. These products were cloned into our previously described single-plasmid reverse genetics system for ZIKV strain MR766 (sequence is available at Genbank accession KX830961)^28^ using techniques like those described in our prior ZIKV envelope and NS2B-NS3 protease protein DMS papers^29,30^. To ensure that we could maintain library complexity at all stages of the screen, we subdivided the NS5 coding sequence into eight 111 to 112 codon segments, hereinafter referred to simply as “tiles.” One added benefit to this tile size was that it fit well into a single paired-end Illumina read, which simplified the deep sequencing and analysis. For each tile, we created three individual libraries to be screened as independent biological replicates.

Libraries were constructed in our previously reported ZIKV MR766 strain cDNA clone^31^, chosen for its high plasmid stability in bacteria, which facilitated efficient cloning, and its ability to generate high-titer viral stocks for robust library production. To avoid the carry-over of replication competent wild type virus as a result of vector contamination, we designed two acceptor plasmids in which either the region corresponding to tile 1 (used for cloning tiles 1-4) or tile 5 (used for cloning tiles 5-8) was replaced by an inert stuffer fragment composed of GFP. We created three mutant plasmid libraries for each tile, which were handled separately in all subsequent steps to provide true biological replicates. Each library contained a minimum of 4×10^5^ unique plasmid clones, which overrepresents the total number of possible codon variants of 3552 (32 combinations of NNS multiplied by 111 codons) by 110-fold, with an average of 1.05 mutations per clone (see Table 4 for summary of each library statistics).

We generated infectious virus stocks of these ZIKV NS5 DMS libraries by transfecting plasmids into HEK 293T cells using a protocol to maintain library complexity as previously reported^32^. Wild-type virus was rescued in parallel as a control. These stocks were titered on Huh-7.5 cells using 4G2 antibody staining and flow cytometry analysis (see below), and then selected by infecting Huh-7.5 cells. Each mutant library was used to infect 2e7 Huh-7.5 cells at a multiplicity of infection (MOI) of 0.05 infectious units per cell, thus maintaining ∼281-fold coverage of each variant. Infected cells were collected at day 2 post-infection. Total RNA was extracted from these cells, reverse transcribed, and subjected to Illumina deep sequencing using a barcoded-subamplicon sequencing approach described in previous work^18^ (see also https://jbloomlab.github.io/dms_tools2/bcsubamp.html) to minimize sequencing errors. The raw deep sequencing data have been deposited in the Sequence Read Archive as BioProject XXXXXXXXXXXX. The code that performs the analyses of the deep sequencing data is available on GitHub at https://github.com/jbloomlab/ZIKV_DMS_NS5_EvansLab. Briefly, we used dms_tools2 (https://jbloomlab.github.io/dms_tools2/)^33^, version 2.4.14, to count the occurrences of each mutation in each sample (see https://jbloomlab.github.io/dms_tools2/bcsubamp.html for details). The amino-acid preferences were computed from these counts using the approach previously described^33^ (see also https://jbloomlab.github.io/dms_tools2/prefs.html). The mutational effects are the log of the preference for the mutant amino acid divided by the preference for the wild-type amino acid. Output containing the numerical values of the counts of mutations in each sample, the amino-acid preferences, and mutational effects are processed on a per-tile basis. These data are provided in CSV file format in the GitHub repository. See the README for details on navigating the analysis output.

### Experimental validation of DMS results

We used a previously described luciferase reporter expressing ZIKV system^29^ to validate the predicted fitness of mutants from the DMS screen. We cloned a subset of 18 individual NS5 mutants with a range of DMS predicted fitness values into a pACYC177 MR766 plasmid that contains a ZIKV MR766 genome under the control of a T7 promoter with a Gaussia luciferase gene inserted between NS1 and NS2A. These plasmids were linearized with ScaI (New England Biolabs, Beverly, MA) and purified with a Zymo DCC-5 column (Zymo Research Corp, Tustin, CA). The linearized DNA was used as a template in a Promega T7 Express In vitro transcription reaction with the addition of m7g cap (Promega, Madison, WI). The reaction was purified with the EZNA RNA Kit (Omega Bio-Tek, Inc., Norcross, GA) and the products were aliquoted to 5 µg per tube and stored in the –80 °C.

For each electroporation, Huh7.5 cells that were washed twice in ice cold PBS were resuspended at 1.5×10^7^ cells/mL, and 400 µL (6×10^6^ cells) were added to 1.5 mL tubes and mixed with 5 µg of RNA. The mixture was transferred to a 1 mm gap cuvette and electroporated a BTX ECM 830 electroporator (BTX Genetronics, Holliston, MA). Cells were rested for 10 minutes at room temperature, then transferred to 10 mL of pre-warmed DMEM with 3% FBS. Electroporated cells were seeded at a density of 1×10^5^ cells per well in a 24-well plate, along with 1×10^5^ non-electroporated cells to act as recipients for spread of virus in the well. Cells were harvested 3 days post electroporation and luciferase activity was measured using a Renilla Luciferase Assay Lysis Buffer (Promega, Madison, WI) and quantified with a BioTex Synergy 4 multidetection microplate reader (BioTek, Winooski, VT).

### Quantification of viral titers

For DMS, the infectious titers of rescued wild type, as a positive control, and mutant libraries were determined by immunostaining with the pan-flavivirus E protein reactive 4G2 antibody and flow cytometry, as previously described^34^. Briefly, one day prior to infection, 24-well plates were seeded at a density of 1 ×10^5^ Huh-7.5 cells/well. The next day, 250 uL of serially diluted transfection supernatants in DMEM with 3% FBS was added to each well (in duplicate). Staining was performed at 24 hours post-infection to avoid any spread. Cells were fixed in 4% paraformaldehyde and stained with 4G2 antibodies and goat-anti-human secondary antibody conjugated to Alexa Fluor 647 (Thermo Fisher Scientific, Waltham, MA) using methods previously described. Data were acquired on an Attune NxY flow cytometer (Thermo Fisher Scientific, Waltham, MA). Only the viral dilutions leading to less than 20% of infected cells were used to calculate the infectious titers. Infectious unit per mL were calculated using the following formula: “percentage of infectious events x number of cells / volume (mL) of viral inoculum”.

Virus titers were also determined by CPE-based limited dilution assay on Huh-7.5 cells, following established protocols^32^. Here, 1×10^4^ cells were seeded in each well of a 96-well plate the day before being infected. For most infections, we used 50 µL of virus serially diluted in DMEM with 10% FBS, 108.8 units/mL penicillin, and 108.8 µg/mL streptomycin (8 wells per dilution). Wells were scored for virus-induced CPE, which was observed as early as 4 days post transfection and infectious titers were calculated as TCID50/mL according to the method of Reed and Muench^35^.

### Multi-cycle growth curves

One day prior to infection, 1×10^5^ Huh-7.5 cells per well were seeded in a 24-well poly-lysine coated plate in triplicate. Then, the cells were infected at a MOI of 0.05 Huh-7.5 cell TCID50 infectious units per cell in DMEM with 3% FBS, 108.8 units/mL penicillin, and 108.8 µg/mL streptomycin. Supernatants were collected daily for the next four days. Supernatant infectivity was assayed by TCID50 assay on STAT2 KO Huh-7.5 cells, to avoid impacts by any IFN produced during the growth curve experiment.

### Mouse experiments

ZIKV stocks were amplified in Vero cells, and cell supernatants were harvested 96 h post infection. Briefly, culture supernatants were collected and virus particles were purified using a 20% sucrose cushion and ultracentrifugation at 24,000 rpm at 4°C for 3h. Viral pellets were then resuspended in PBS. Virus stocks were titrated using a plaque forming unit (PFU) assay on Vero cells and aliquoted were stored at −80°C. For PFU assays, virus sample was serially diluted in RPMI complete media and then added to Vero cell monolayers in 24 well plates. After incubation at 37C for 1 hour, cells were overlaid with 0.8 % methylcellulose. After 5 days, cells were fixed with 4% paraformaldehyde and stained with crystal violet for visualization of plaques.

Mouse studies were carried out in an animal biosafety level 2 facility under a protocol approved by the Icahn School of Medicine at Mount Sinai Animal Care and Use Committee. Human STAT2^KI/KI^ (*hSTAT2^KI/KI^*) mice were injected intravenously with 100 μl of PBS (mock), wild type (WT) or double mutant virus at a dose of 1×10^6^ PFU per mouse. Stat2^-/-^ (*Stat2^-/-^*) mice were injected subcutaneously with 100 μl PBS, WT or double mutant virus at a dose of 1×10^4^ PFU per mouse. Mice were monitored daily for weight loss, symptom development, and survival for 18 days. Symptom development was measured based on a 5-point scale system: 1.0, ruffled fur; 2.0, lethargy; 3.0, hunched back and limited mobility; 4.0, moribund/ paralysis; 5.0 found dead. After 18-days, Kaplan–Meier survival graphs were plotted and differences were analysed by Long-rank (Mantel-Cox) test.

### Statistical analysis

Data were analyzed using an unpaired t-test with Welch’s correction, either with or without a correction for multiple comparisons. Asterisks indicate the following P-value significance. *P<0.05, **P<0.005, ***P<0.0005 on Prism software (GraphPad Software). Values in graphs represent the mean and standard error of experiments performed in triplicate or quadruplicate, with between two and five independent experiments.

## Acknowledgements

This study was supported in part by NIH grants R01 AI175303 (M.J.E. and A.G.S.), R01 AI166594 (JKL and MJE) and F31 FAI191695A (EB). The authors thank Charles Rice (Rockefeller University, New York, NY) for 293T and Huh-7.5 cells, Paul Bieniasz (Rockefeller University, New York City, NY) for kindly providing the pSCRPSY vector, and Florian Krammer for 4G2 monoclonal antibody (Icahn School of Medicine at Mount Sinai, New York, NY). We would like to thank the expertise and assistance of the Dean’s Flow Cytometry CORE at Mount Sinai. The authors want to thank Charles Rice (Rockefeller University, New York, NY) for kindly providing Huh-7.5 cells and Paul Bieniasz (Rockefeller University, New York City, NY) for kindly providing the pSCRPSY vector. JDB is an Investigator of the Howard Hughes Medical Institute. The funders had no role in study design, data collection and analysis, decision to publish, or preparation of the manuscript.

## Author Contributions

R.B.R. and M.J.E. conceived this study. R.B.R., C.K., A.G., E.B., M.S.S., B.D., E.C.V., A.H. collected, analyzed, and interpreted data. A. G.-S., J.D.B., J.K.L., and M.J.E. supervised this study. A.G.S., J.K.L., and M.J.E. acquired funding for the project. R.B.R. and M.J.E. wrote the manuscript. All authors reviewed the results and approved the final version of the manuscript.

## Data and code availability

See the GitHub repository https://github.com/jbloomlab/ZIKV_DMS_NS5_EvansLab for all analysis and code, including:

- The quality checks on the deep mutational scanning data across all library tiles, including summaries of read depth, mutation frequency, comparisons of plasmid and virus libraries, and other key metrics of library quality https://github.com/jbloomlab/ZIKV_DMS_NS5_EvansLab/tree/main/results/notebooks
- The mutation effects across all tiles of the library in CSV format https://github.com/jbloomlab/ZIKV_DMS_NS5_EvansLab/blob/main/results/all_tiles/alltiles_host_adaptation.csv
- The PDB structure used for viewing deep mutational scanning data projected onto protein https://github.com/jbloomlab/ZIKV_DMS_NS5_EvansLab/blob/main/data/NS5_STAT2_joined.pdb

Source data are provided with this paper.

---

**Supplementary Fig. 1:**
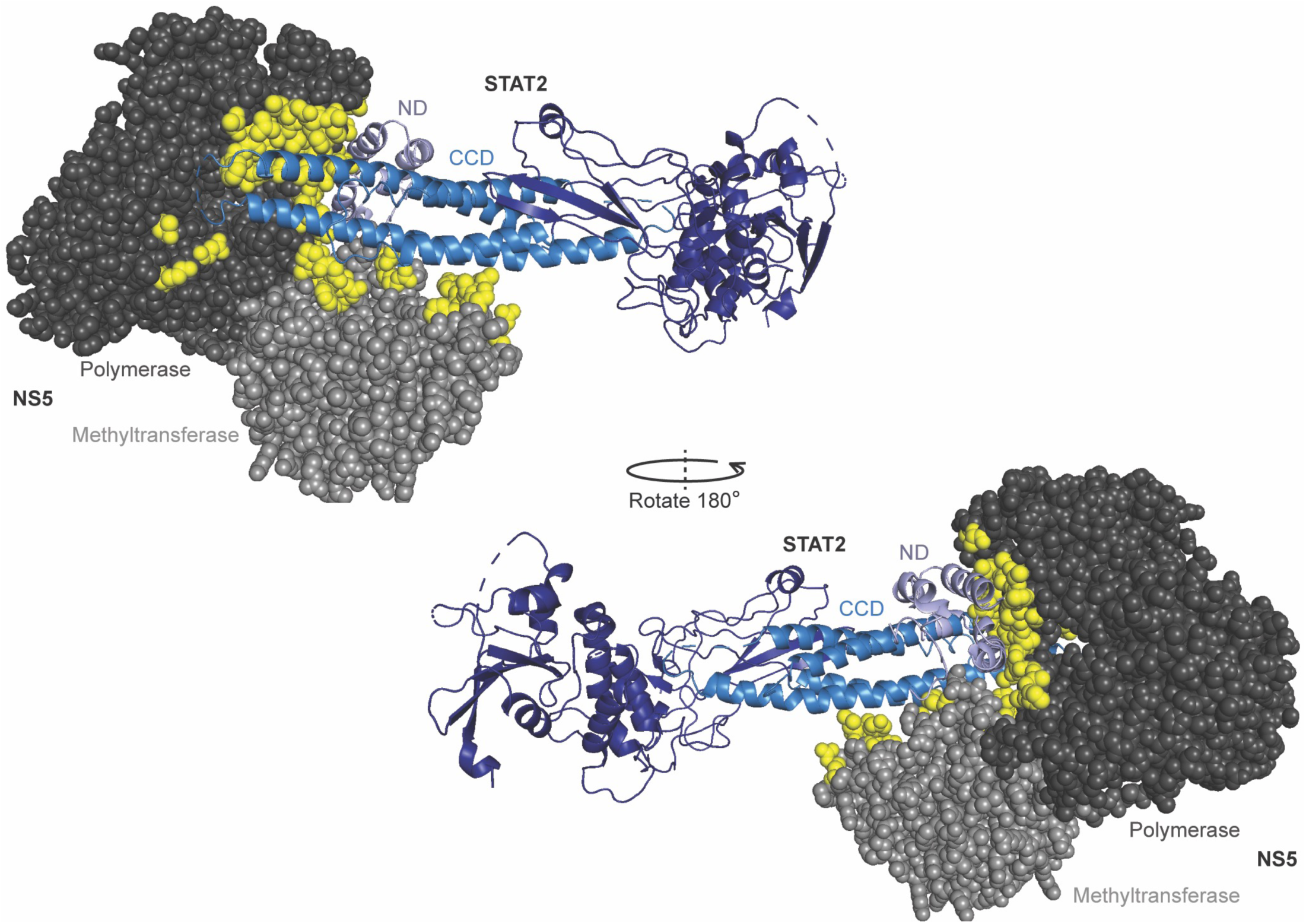
Structural model of the ZIKV NS5–STAT2 complex. Composite structural model showing the interaction between ZIKV NS5 and human STAT2. NS5 is displayed as a space-filling model with the methyltransferase (light gray) and RNA-dependent RNA polymerase (dark gray) domains indicated. STAT2 is shown as a ribbon diagram, with the N-terminal domain (ND) in light blue and the coiled-coil domain (CCD) in marine blue. NS5 residues that lie in close proximity defined to STAT2 are highlighted in yellow. Two orientations of the complex are shown, rotated 180° around the vertical axis to reveal both sides of the interface. Structures were modeled using available cryo-EM and crystallographic data (PDB: 6WCZ, 6UX2).

**Supplementary Fig. 2:**
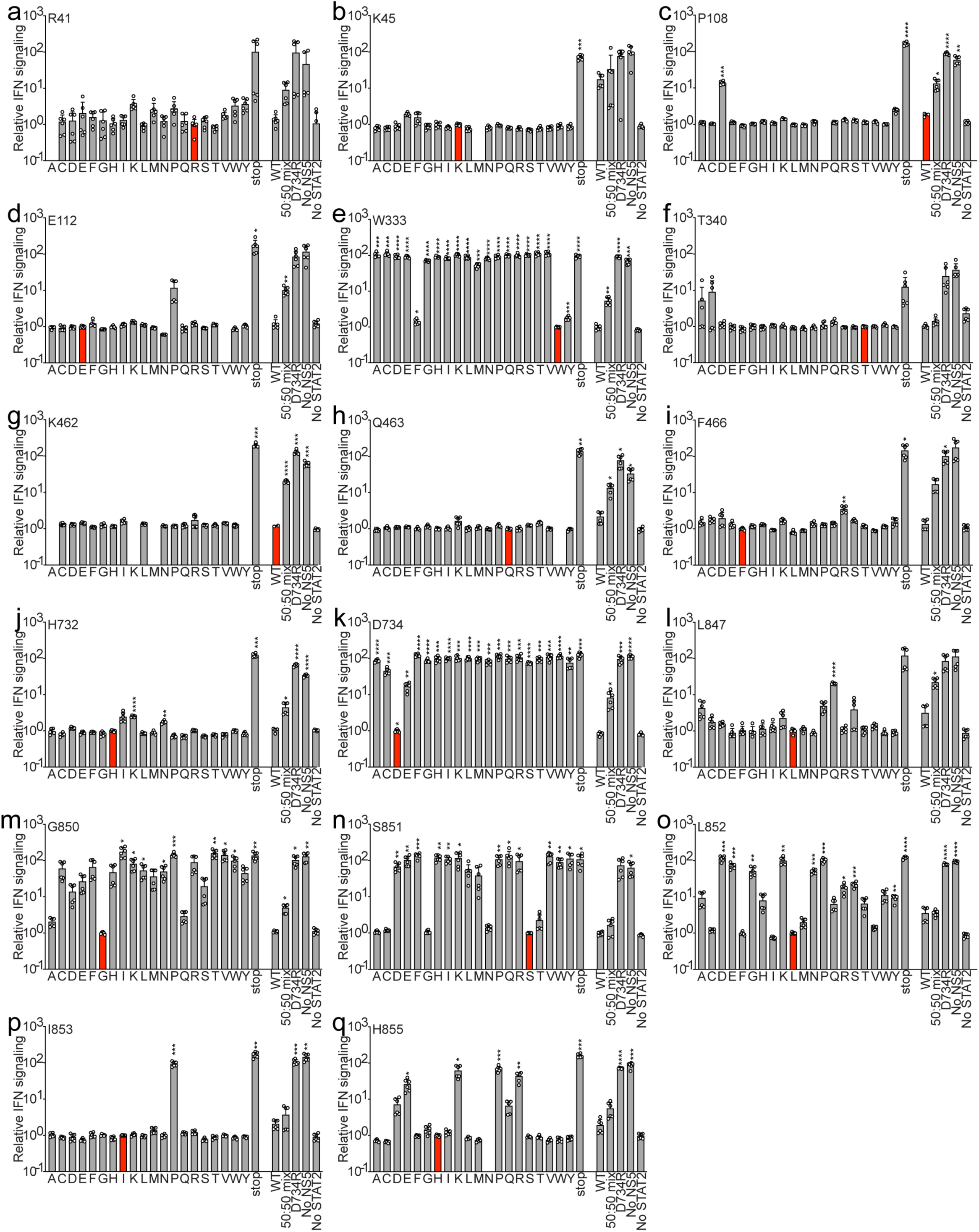
Impacts of single amino acid mutations on ZIKV NS5 IFN antagonism. Each panel shows the relative IFN signaling in STAT2 knockout 293T cells in the presence of single amino acid substitutions at the residue indicated on the x-axis. Each panel includes the controls: wild type NS5, the antagonism deficient D734R mutant, an equal mixture of wild type and D734R, ‘no NS5’, and ‘no NS5 or STAT2’. All transfections except the ‘no NS5 or STAT2’ condition include human STAT2 to complement the KO. Asterisks indicate statistical significance determined by multiple unpaired t-tests with Holm-Šidák method to correct for multiple comparisons.

**Supplementary Fig. 3:**
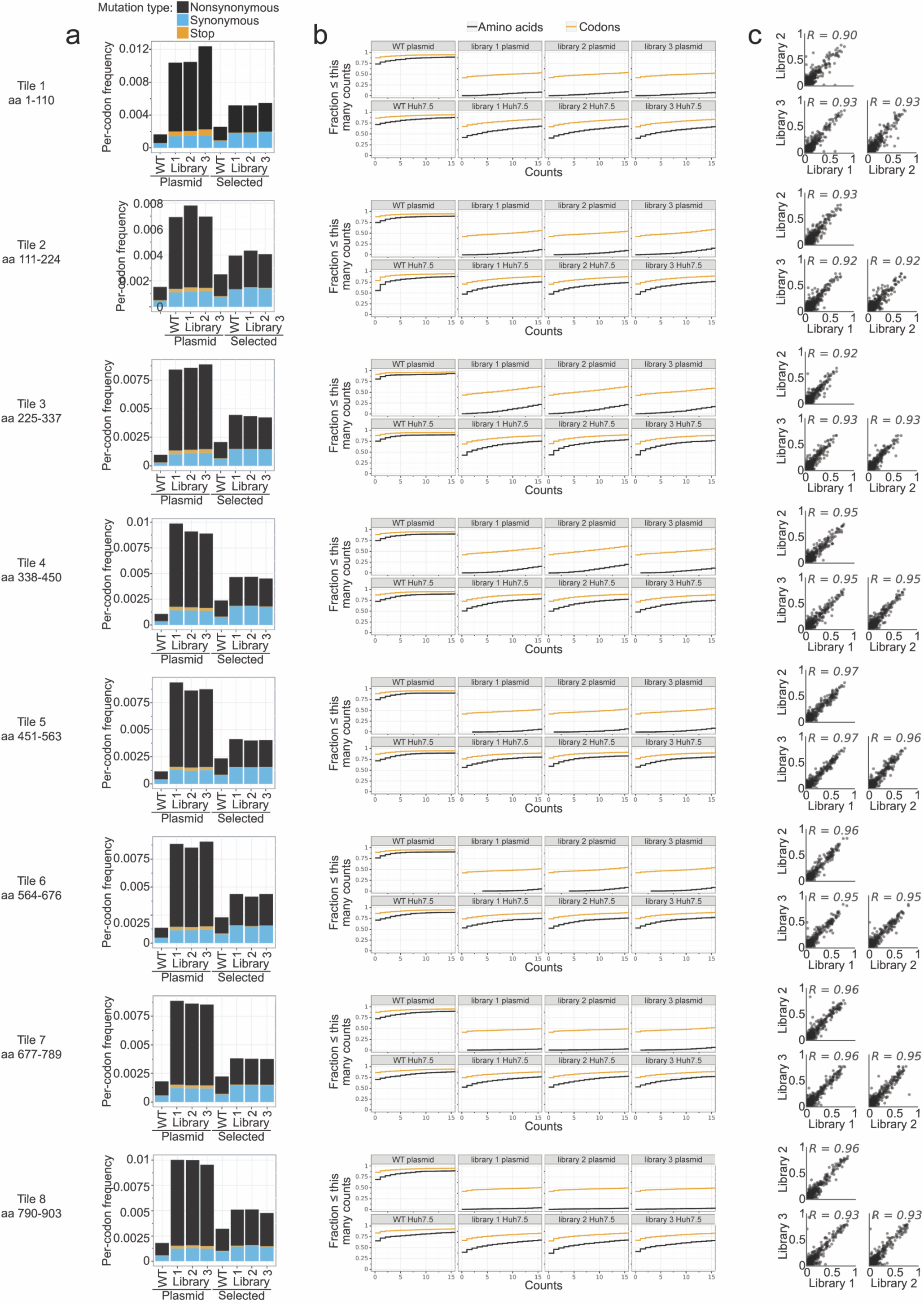
Quality control and validation of ZIKV NS5 deep mutational scanning libraries. The NS5 coding region was divided into eight consecutive ‘tiles’ for DMS. Tile numbers and amino-acid ranges are labeled to the left of each row, with the remaining panels in each row showing: **(a)** Per-codon mutation frequencies measured in plasmid DNA (input) and after viral selection are shown, with mutation types indicated (nonsynonymous = black, synonymous = blue, stop = orange). **(b)** Cumulative count distributions for amino acid substitutions (black) and codons (orange) are displayed for plasmid libraries and post-selection viral samples. **(c)** Scatterplots show pairwise correlations of codon frequencies between independently generated plasmid libraries, with Pearson correlation coefficients (R) indicated.

